# State-Dependent Lipid Interactions with the A2a Receptor Revealed by MD Simulations Using *In Vivo*-Mimetic Membranes

**DOI:** 10.1101/362970

**Authors:** Wanling Song, Hsin-Yung Yen, Carol V. Robinson, Mark S.P. Sansom

## Abstract

G protein-coupled receptors (GPCRs) are the largest family of integral membrane proteins and a major class of drug targets. Membranes are known to have modulatory effects on GPCRs via specific lipid interactions. However, the mechanisms of such modulations in cell membranes and how they influence GPCR functions remain unclear. Here we report coarse-grained MD simulations on the Adenosine A2a receptor embedded in an in vivo mimetic membrane model comprised of 10 different lipid species. Three conformational states of the receptor, i.e. the inactive state, the active state, and the active state with a mini-G_S_ protein bound were simulated to study the impact of protein-lipid interactions on the receptor activation. The simulations revealed three specific lipids (GM3, cholesterol and PIP_2_) that form stable and preferential interactions with the receptor, differentiating these from bulk lipids such as PS, PE and PC. In total, nine specific lipid-binding sites were revealed. The strength of lipid interaction with these sites depends on the conformational state of the receptor, suggesting that these lipids may regulate the conformational dynamics of the receptor. In particular, we revealed a dual role of PIP_2_ in promoting A2aR activation, which involves stabilization of both the characteristic outward tilt of helix TM6 within receptor and also the association of A2aR and mini-Gs when the activated complex forms. Structural comparisons suggested that PIP_2_ may facilitate Gα activation. Our results reveal likely allosteric effects of bound lipids in regulating the functional behaviour of GPCRs, providing a springboard for design of allosteric modulators of these biomedically important receptors.

---

The G protein-coupled receptors (GPCRs) form the largest super-family in the mammalian genome. They bind to a wide range of ligands and convert extracellular signals to intracellular responses via interactions with either G proteins or β-arrestins (Zhou et al., 2017). Owing to their involvement in many physiological processes, GPCRs are targeted by about 30% of current drugs. Recent advances in membrane protein structural biology have greatly expanded our understanding of GPCRs. GPCRs share a conserved architecture of 7 transmembrane helices (TM1-7) connected by three extracellular loops (ECL1-3) and three intracellular loops (ICL1-3). The structures of four Class A GPCRs (rhodopsin(Choe et al., 2011; Kang et al., 2015; Scheerer et al., 2008), the β2 adrenergic receptor (Rasmussen et al., 2011a; Rasmussen et al., 2011b; Ring et al., 2013), M_2_ muscarinic receptor (Kruse et al., 2013) and A2a receptors (Carpenter et al., 2016)), and of two Class B GPCRs (the CT receptor (Liang et al., 2017) and the GLP-1 receptor (Liang et al., 2018; Zhang et al., 2017)) have been determined in active states stabilized by auxiliary proteins of G proteins, β-arrestin or antibodies. Comparison between the inactive and active state structures have revealed the conformational changes during receptor activation, which include the opening of an intracellular binding pocket that is achieved by a large outward pivotal tilt of TM6 accompanied by movements of the TM5, TM4 and TM3 helices, and also include adjustments in the ligand binding pocket and extracellular loops brought about by bound ligands. The sharp contrast between the relatively conserved orthosteric binding pockets and the wide spectrum of signals which GPCRs are able to elicit has resulted in a search for allosteric modulators that could fine-tune the conformational dynamics of the receptors. Several recent structures have revealed allosteric binding sites on the extra-helical surface of GPCRs (Jazayeri et al., 2016; Song et al., 2017; Zhang et al., 2015), emphasizing the potential for modulation from outside of the helix bundles.

Lipid bilayer membranes have been shown to modulate various GPCR activities, including ligand binding, conformation stability and oligomerization (Oates and Watts, 2011). Modulatory effects mediated via changes in membrane physical properties, e.g. membrane thickness, curvature and surface tension, have been studied extensively by both experimental and computational methods (Chachisvilis et al., 2006; Mondal et al., 2013; Periole et al., 2007). Recently, modulation of GPCRs via specific interactions with lipids have gained attention. Thus, phosphatidylglycerol (PG) modulates the interaction between a G protein and the neurotensin receptor NTS1 (Inagaki et al., 2012); different lipid headgroup types are able to stabilize different conformational states of the β2 adrenergic receptor (Dawaliby et al., 2015); and anionic lipids such as PIP_2_ facilitate functional interaction between the β2 adrenergic receptor and GRK5 (Komolov et al., 2017).

Molecular dynamics (MD) simulations have provided structural insights into GPCR-lipid interactions. Atomistic simulations identified multiple cholesterol binding sites on the surface of GPCRs, the occupancy of which resulted in increased conformational stability (Guixa-Gonzalez et al., 2017; Lyman et al., 2009; Manna et al., 2016). Frequent insertion of PG into the opening between TM6 and TM7 was observed in atomistic simulations of β2 adrenergic receptor in the active state conformation, suggesting a possible explanation for the influence of anionic lipids on GPCR activation (Neale et al., 2015). Coarse-grained (CG) methods (using e.g. the Martini model (Marrink et al., 2007; Monticelli et al., 2008)) allow for simulation of extended duration which sample more efficiently the diffusion of lipids, providing an unbiased picture of the interactions of lipids with integral membrane proteins (Corradi et al., 2017). Thus CG simulations have revealed that the binding of cholesterol to GPCRs is dependent on cholesterol concentration and influences dimerization kinetics and the resultant dimer interfaces (Pluhackova et al., 2016; Prasanna et al., 2014; Prasanna et al., 2016; Provasi et al., 2015). Recent CG simulations using bilayers comprised of multiple lipid species have provided insights into GPCR-lipid interactions in a more biologically realistic environment (Ingolfsson et al., 2014; Koldso and Sansom, 2015). For example, the μ-opioid receptor embedded in a complex lipid membrane was shown to induce lipid regions with high-order near certain transmembrane helices that may facilitate receptor dimerization (Marino et al., 2016). However, it remains unclear that how GPCR-lipid interactions modulate the functions of GPCRs such as receptor activation and downstream signalling in a physiologically relevant context, i.e. in a lipid bilayer environment mimicking a mammalian cell membrane.

In this study, we employ CG MD simulations to characterise the interactions of lipids with GPCRs in complex in vivo-mimetic membranes. We focus on the Adenosine A2a receptor, a prototypical GPCR that plays a major role in central nerve system in response to adenosine, as its structure has been determined in both an inactive (Jaakola et al., 2008) and active (Carpenter et al., 2016) state. The active state of the receptor was determined in complex with an agonist and an engineered G protein (‘mini Gs’) which binds to the activated receptor in a conformation virtually identical to that observed in the β2AR-Gs structure (Carpenter et al., 2016). Comparing the protein-lipid interactions in three conformational states, i.e. the inactive state, the active state, and the active + mini Gs state, we have characterised the interactions of 10 physiologically relevant lipid species with the receptor and changes of these interactions in response to receptor activation. We observed a clear distinction between those lipids that form specific interactions with the receptor (namely GM3, cholesterol and PIP_2_) and the remaining bulk lipids (namely PC, PE, PS and sphingomyelin species). The strength of specific lipid interactions with the A2aR showed a degree of sensitivity to the conformational state of the receptor, suggesting that these lipids may play a role in regulating its conformational dynamics. At the intracellular side of A2aR, we observed four PIP_2_ binding sites that are conserved across Class A GPCRs. Potential of mean force (PMF) calculations of the free energy landscape of GPCR/PIP_2_ and GPCR/mini Gs interactions suggest that bound PIP_2_ molecules may have dual functional effects on both receptor activation and enhancing A2a-mini Gs association. Our results suggest that lipid interaction sites may provide new targets for drugs acting as allosteric modulators of GPCRs.

## RESULTS

### GM3, Cholesterol and PIP2 interact with the A2aR

To explore the possible modulatory role of membrane lipids on A2a receptor activation, we performed CG MD simulations of the receptor in three different conformations, namely an inactive state, an active state, and an active state with bound mini Gs protein (see Figure 1A and SI Table S3). The simulations were of single copy of the receptor in an asymmetric lipid bilayer comprised of 10 different lipid species providing an *in vivo*-mimetic membrane environment (Figure 1B).

**Figure 1.**
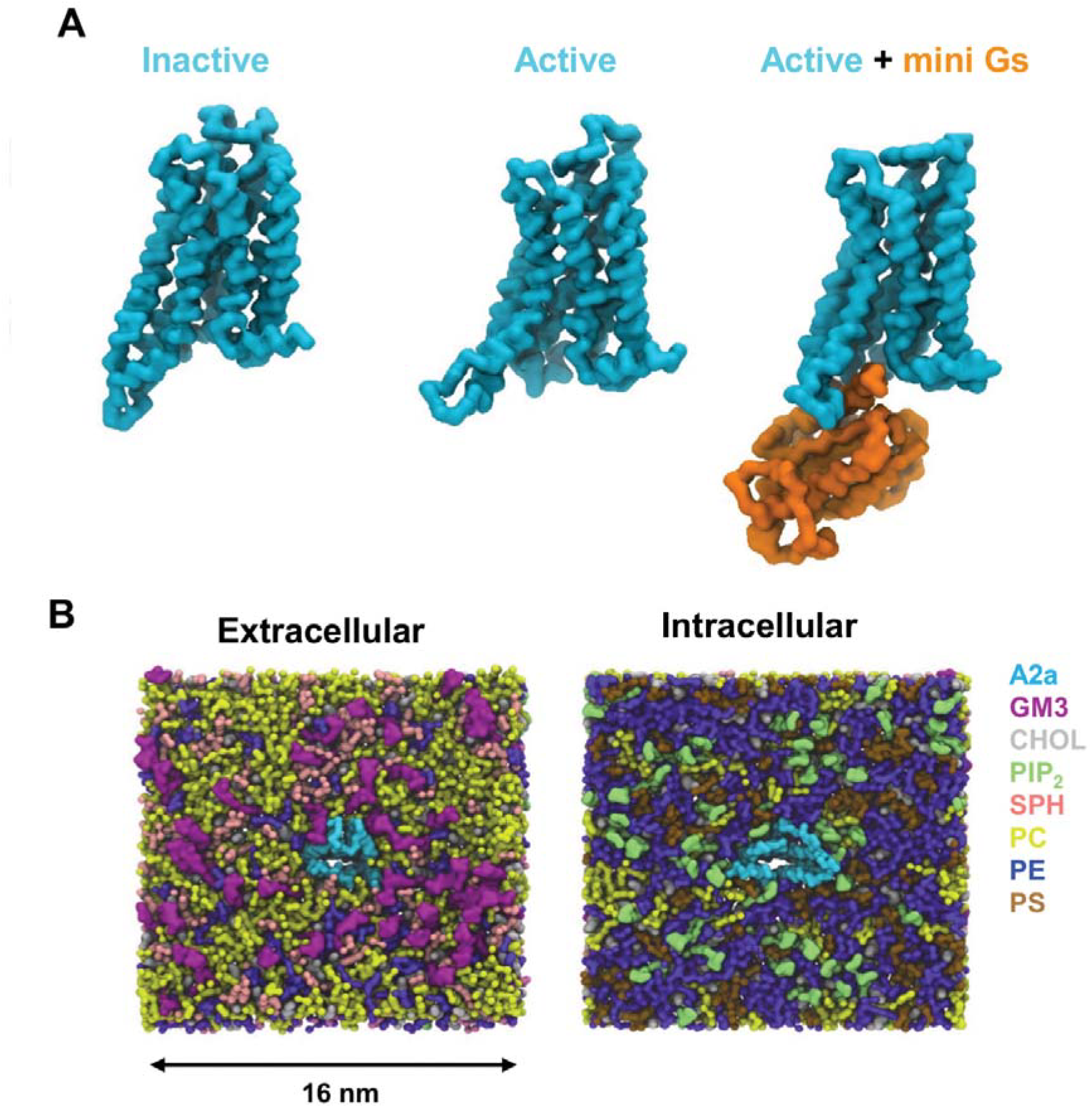
**A** Three different conformational states (inactive, PDB code 3EML; active, PDB code 5G53, subunit A, and active + miniGs, PDB code 3G53, subunits A and C) of the A2aR used in the simulations. **B** An overview of the simulation system from the extracellular and intracellular sides. The receptor is coloured cyan and different lipid species are coloured as specified. In the extracellular view, GM3 is in purple and in the intracellular view, PIP_2_ is in green.

Analysis of the area per lipid (APL) as a function of time (SI Figure S1) showed that the simulation systems did not exhibit abrupt or significant deformation during the course of the simulations. Average APLs (SI Table S5) indicate that the upper leaflet is somewhat better ordered and more tightly packed than the lower leaflet, largely due to the lower degree of tail unsaturation in the upper leaflet. Cholesterol, which initially was present in equal concentrations in the two leaflets, accumulated to the outer leaflet to a small degree, due to its preference for interaction with saturated lipid tails. The APLs of cholesterol between the two leaflets, however showed no significant difference.

Radial distribution functions (RDFs; Figure 2A) revealed the 10 species of lipids could be divided into two groups based on their proximity to the surface of the receptor in all three conformational states: Group 1 formed close contacts with the receptor and included GM3, cholesterol and PIP_2_; Group 2 lipids (referred to from now on as bulk lipids) did not form frequent close interactions with the receptor and included PC, PE, PS species and sphingomyelin. Monitoring the number of lipids of each species within the first shell around the receptor (defined as within 1 nm of the receptor surface as indicated by RDFs) showed that the exchange between the first shell and bulk lipids reached equilibrium at ~3 μs (SI Figure S2). Consequently, the protein-lipid interaction analyses in this paper were based on data collected from the period 3-8 μs. At equilibrium, the receptor in the inactive state, the active state, and the active + mini Gs state were surrounded by 13 ± 2 (average value ± standard deviation), 16 ± 2 and 17 ± 3 PIP_2_ molecules in the lower leaflet and 14 ± 2, 13 ±3, 13 ±3 GM3 molecules in the upper leaflet respectively. Cholesterol showed an asymmetric distribution around the receptor in the two leaflets. Thus, the receptor in the inactive state, the active state, and the active + mini Gs state was surrounded by 7 ± 2, 8 ± 2, 8 ± 2 cholesterol molecules in the upper leaflet and 13 ± 2, 13 ± 2, 13 ± 3 cholesterol molecules in the lower leaflet respectively.

**Figure 2.**
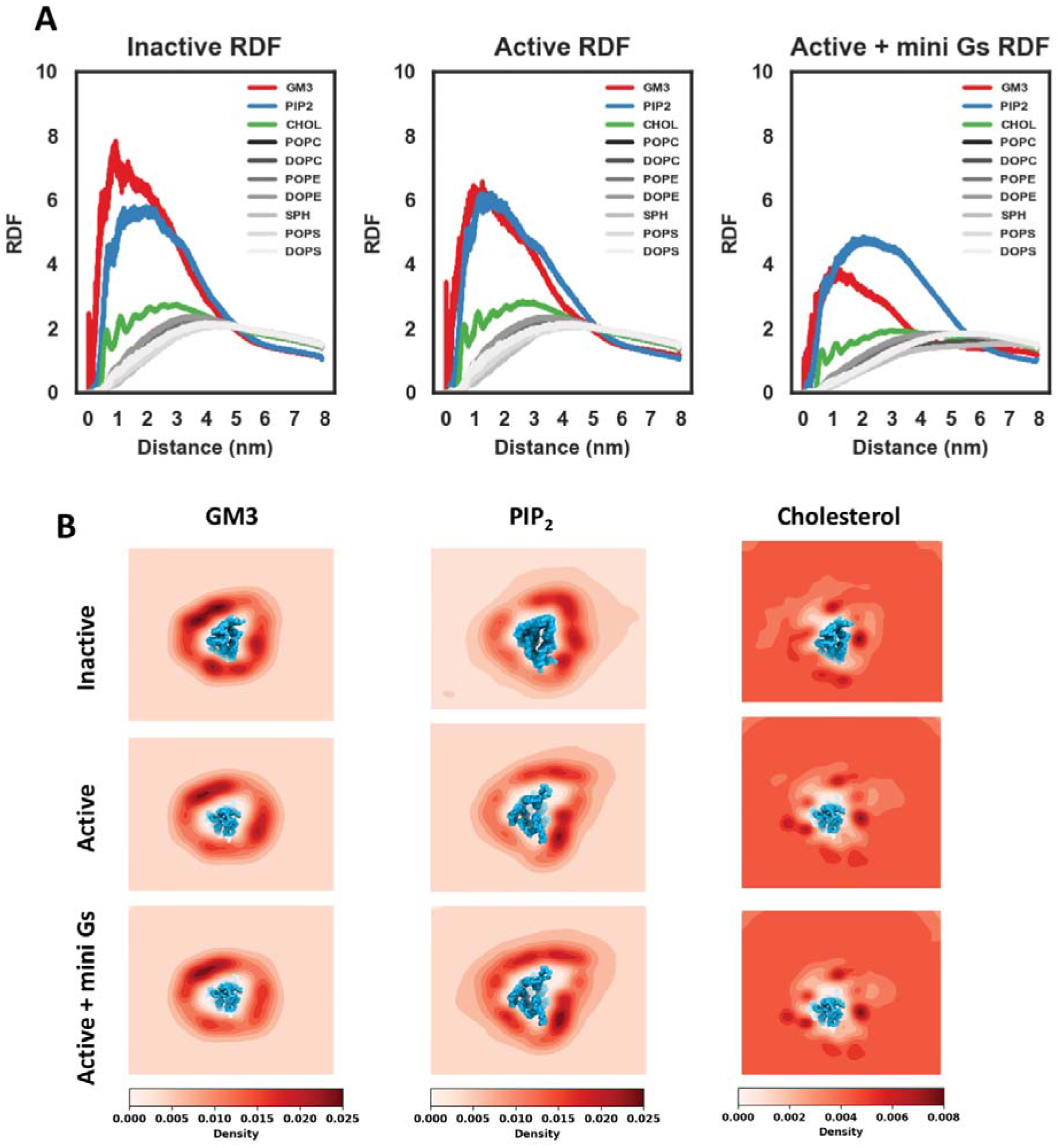
**A** Radial distribution functions of lipid around the protein for the three conformational states of the receptor. The RDFs of lipids with specific interactions are colour coded while those of the bulk lipids are in grey shades. The RDFs were averaged over the 10 simulations of each conformational state (see SI Table S3). **B** Density of GM3, cholesterols and PIP_2_ around the receptor in different conformational states. The density maps were averaged over the 10 simulations of each conformational state.

The 2D density in the membrane plane was calculated for each lipid species to check their binding locations on the receptor surface. According to the density maps, the bulk lipids showed layers of lipid shells surrounding the receptor in all states with no specific binding site (SI Figure S3 and S4). In contrast, strongly preferred binding locations were clearly observed for cholesterol, GM3 and PIP_2_ (Figure 2B). The binding locations of cholesterols, GM3 and PIP_2_ did not vary much when comparing between different conformational states of the receptor, with preferred binding locations at TM1-4, TM4/ECL2/TM5, and TM6/TM7 for GM3; TM1-TM2, TM3/TM5, TM5-TM6 and TM7/H8 for PIP_2_; and TM1/TM2, TM3/TM5, TM6/TM7 and multiple binding spots surrounding TM4 for cholesterols. However, the relative binding probabilities at these locations were clearly dependent on the state of the receptor, indicating that the binding affinities of these locations are sensitive to the receptor activation state.

### Nine lipid binding sites revealed by simulations

To identify the specific binding sites for each lipid species, we calculated the interaction duration per residue, i.e. the average duration of the continuous contacts between a given lipid species and the residue. Based on this measurement, we were able to identify 9 distinct lipid binding sites on the surface of A2a receptor (SI Table S7). Together, these account for nearly all the hydrophobic grooves on the transmembrane surface of the receptor. These binding sites were conserved across the three conformational states and were pre-dominantly occupied by one or two lipid species from Group 1 (i.e. GM3, cholesterol or PIP_2_) whilst remaining accessible to lipids from the bulk (Figure 3 and SI Figure S5). The distribution of interaction durations of Group 1 lipids with the identified binding sites were fitted as mono-exponential decay curves. We therefore calculated the *k*_*off*_ values of lipids from the 9 identified binding sites from the decay of interaction durations of the residues that showed the strongest interactions with the given species of lipid within their binding sites (Table 1 and SI Figure S6). The *k*_*off*_ values of GM3, cholesterols and PIP_2_ ranged from 2 to 14 μs^−1^. This together with an average number of binding events of 500-2500 to the residues in the interaction sites is indicative of sufficient sampling of both binding locations and binding poses in our simulations.

**Figure 3.**
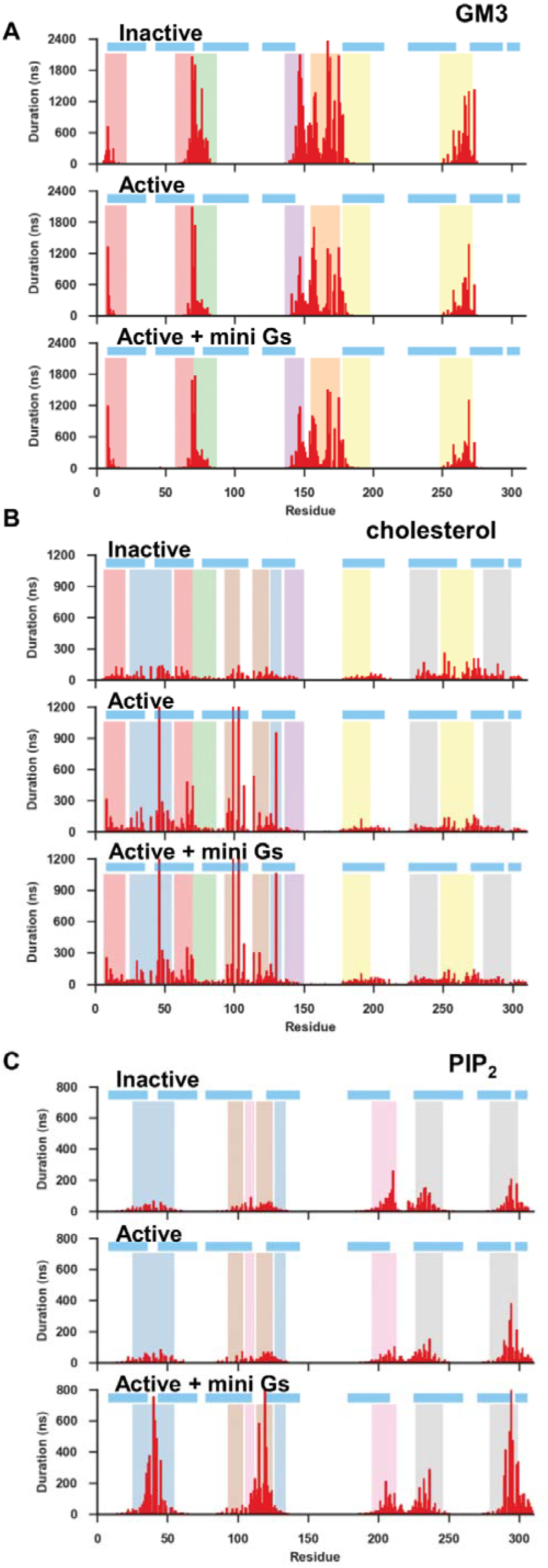
Average duration of lipid interactions with the three states of the receptor as a function of residue number for GM3 (**A**), PIP_2_ (**B**) and cholesterol (**C**). The horizontal blue lines indicate the positions of the transmembrane helices, and the vertical coloured bands indicate the 9 lipid binding sites identified from this analysis (also see Fig. 4).

**Table 1.** *k*_*off*_ of Group 1 lipids dissociating from the nine identified binding sites.

| Binding Sites | $k_{off}(\mu s^{-1})$ | | |
| --- | --- | --- | --- |
|  | Inactive | Active | Active + mini Gs |
| Nter/TM1/TM2 | 11 (CHOL)<br>4 (GM3) | 9 (CHOL)<br>4 (GM3) | 8 (CHOL)<br>3 (GM3) |
| TM1/TM2/TM4 | 7 (CHOL)<br>7 (PIP<sub>2</sub>) | 4 (CHOL)<br>6 (PIP<sub>2</sub>) | 4 (CHOL)<br>3 (PIP<sub>2</sub>) |
| TM2/ECL1/TM3 | 11 (CHOL)<br>2 (GM3) | 9 (CHOL)<br>3 (GM3) | 8 (CHOL)<br>5 (GM3) |
| TM3/TM4/ECL2 | 8 (CHOL)<br>4 (GM3) | 13 (CHOL)<br>4 (GM3) | 10 (CHOL)<br>5 (GM3) |
| TM3/ICL2/TM4 | 7 (CHOL)<br>6 (PIP<sub>2</sub>) | 7 (CHOL)<br>4 (PIP<sub>2</sub>) | 6 (CHOL)<br>3 (PIP<sub>2</sub>) |
| TM3/TM5 | 5 (PIP<sub>2</sub>) | 3 (PIP<sub>2</sub>) | 5 (PIP<sub>2</sub>) |
| TM4/ECL2/TM5 | 4 (GM3) | 4 (GM3) | 5 (GM3) |
| TM5/ECL3/TM6 | 8 (CHOL)<br>7 (GM3) | 8 (CHOL)<br>5 (GM3) | 7 (CHOL)<br>6 (GM3) |
| TM6/TM7 | 9 (CHOL)<br>4 (PIP<sub>2</sub>) | 7 (CHOL)<br>2 (PIP<sub>2</sub>) | 7 (CHOL)<br>3 (PIP<sub>2</sub>) |

#### GM3

GM3 lipid molecules exhibited five interaction sites on A2aR at: (i) the N-terminus/TM 1/TM2; (ii) TM2/ECL1/TM3; (iii) TM3/TM4/ECL2; (iv) TM4/ECL2/TM5, and (v) TM5/ECL3/TM6 (Figures 3A and 4A). A conserved binding mode was revealed: the N-acetyl neuraminic acid (Neu5Ac) moiety of the lipid head group (corresponding to the GM13 bead in Martini CG model) interacted with Asn and Gln sidechains on the extracellular loops, and the sugar rings and the lipid tails stacked against adjacent Trp/Leu/Ile residues (Figure 4B). State dependent differences in GM3 interaction durations were observed at: (i) the N-terminus/TM1/TM2, with an increase in mean duration of interaction from ~0.6 μs for the inactive state to ~1.2 μs for the active state; (ii) TM3/TM4/ECL2, a decrease from ~2 μs in the inactive state to ~1 μs in the active state; and (iii) TM4/ECL2/TM5, a decrease from ~2.4 μs in the inactive state to ~1.2 μs in the active state. The former increase in duration at the N-terminus/TM1/TM2 was due to a local unwinding of TM2 above the kink at G56^2.54^ (where the superscripts refer to Ballesteros–Weinstein numbering (Ballesteros and Weinstein)) in the active conformation that increases the inter-helix distance between TM1 and TM2 and consequently increases the hydrophobic contact between the receptor and the lipid tail. The latter two decreases were due to the conformational changes in the ECL2 and shifts along their corresponding helical axes of the extracellular end of TM3, TM4 and TM5 that resulted from the agonist-induced binding pocket shrinkage (Carpenter et al., 2016; Lebon et al., 2011). The closer inter-helical distance thus decreased the binding pocket volume for GM3 in the active conformation.

**Figure 4.**
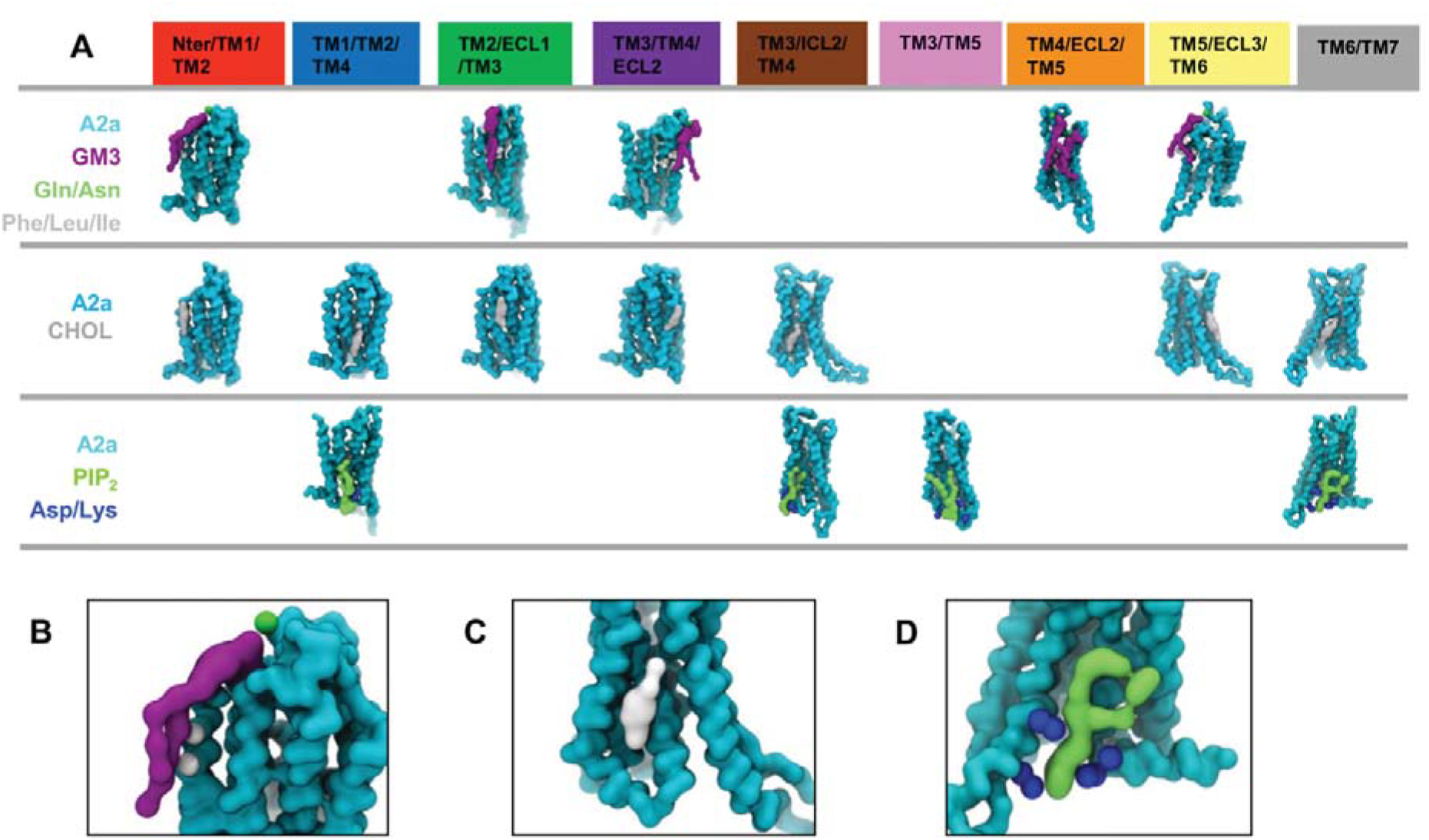
(**A**) Representative binding poses of GM3, cholesterol, and PIP_2_ for each of the binding sites identified by the analysis of lipid interactions shown in Fig. 3. Zoomed in images are provided for examples of these interactions: the A2aR/GM3 (purple) interaction at the Nter/TM1/TM2 site (**B**); the A2aR/cholesterol (grey) interaction at the TM3/ICL2/TM4 site (**C**); and the A2aR/PIP_2_ interaction at the TM6/TM7 site (**D**).

#### Cholesterol

Our simulations revealed seven cholesterol interaction sites, covering nearly all the hydrophobic grooves between the transmembrane helices of the A2aR (Figures 3B and 4A). Cholesterol, and its more water soluble analogue cholesteryl hemisuccinate, are frequently used in crystallization to enhance the thermo-stability of proteins. The available crystal structures of GPCRs to date have revealed 8 cholesterol binding sites (SI Table S6), four of which (i.e. TM2/ECL1/TM3, TM1/TM2/TM4, TM3/ICL2/TM4 and TM6/TM7) demonstrated stable binding in our simulations, while the other 4 showed cholesterol interactions albeit with lower stability.

The duration of interactions of cholesterol molecules with A2aR seen in our simulations showed a degree of dependence on the conformational state of the receptor. This resulted from both the conformational state of the receptor and from interplay with other lipid species binding at the same or overlapping sites (see SI Table S7). For instance, the interaction duration of cholesterol molecules at sites TM1/TM2/TM4 and TM3/ECL2/TM4 were significantly increased in the receptor active and active + mini Gs states. This reflected both the shift of TM4 along its helical axis in the active conformation that led to tighter interactions of the receptor with cholesterols at these two sites, and also the tighter binding of PIP2 at these sites (see below) that blocked the exit routes of cholesterols (Figure 4A). Similar synergistic interplays were observed between GM3 and cholesterol at the binding site defined by Nter/TM1/TM2 where both lipids showed increased interaction duration in the active and active + mini Gs states. Competing interactions were also observed. For example, the interaction duration of cholesterols at site TM6/TM7 was decreased in the active and active + mini Gs states, because PIP_2_ displaced bound cholesterol from the site by binding deep into the opening between TM6 and TM7 in the active state (see below).

#### PIP_2_

Four PIP_2_ binding sites were revealed by our simulations, at the intracellular rim of the receptor adjacent to: (i) TM1/TM2/TM4; (ii) TM3/ICL2/TM4; (iii) TM3/TM5; and (iv) TM6/TM7 (Figures 3C and 4A). The PIP_2_ molecules bound to these sites via interactions between the polyanionic phosphorylated inositol headgroup and basic residues in the binding sites, i.e. R107^3.55^, R111^34.52^, R120^4.41^, K122^4.43^, R205^5.66^, R206^5.67^, K227^6.29^, K233^6.35^, R291^7.56^, and R293^8.48^. Structure-based sequence alignment of the available Class A GPCR structures revealed that these identified basic residues at the intracellular side of the receptors are conserved, and hence the four PIP_2_ binding sites may be common features across the Class A GPCRs (Yen et al., 2018). We also note that the interaction of PIP2 with GPCRs is unlikely to be driven solely by electrostatic interactions, as recent mass spectrometry experiments on β1AR have revealed a significantly lower binding affinity of PIP_3_ (Yen et al., 2018). Comparison between PIP_2_ interactions with conformational states revealed that the interaction duration of PIP_2_ at binding sites TM1/TM2/TM4, TM3/TM5, and TM3/ICL2/TM4 was increased when mini Gs was in complex with the receptor. For the TM1/TM2/TM4 and TM3/TM5 sites the duration increased from ~100 ns in the inactive and active states to ~ 800 ns in the active + mini Gs state, whereas the latter one showed a shift of contacts from R206^5.67^ and K209^5.70^ on TM5 to R107^355^ and R111^34.52^ on TM3 and ICL2. This shift of interacting fingerprints led to the interaction duration at TM3 increased to ~220 ns in the active + mini Gs state from ~50 ns in the inactive or active state. For the binding site TM6/TM7, the PIP_2_ interaction duration was increased to ~400 ns in the active state from ~200 ns in the inactive state, and further increase to ~800 ns in the active + mini Gs state.

### The Energetics of PIP_2_ Interactions

To understand in more detail the relationship of changes in PIP_2_ binding to receptor activation and mini Gs association, we calculated potential of mean forces (PMFs) for the interactions of PIP_2_ with the binding sites identified on the receptor. PMF calculations using coarse-grained MD simulations have been applied to study the energetics of protein-lipid interactions for a number of membrane proteins, including mitochondrial respiratory chain complexes (Arnarez et al., 2013) and transporters (Hedger et al., 2016a), ion channels (Domanski et al., 2017), and EGF receptors (Hedger et al., 2016b). These calculations have been shown to reveal the strength and specificity of the interactions of anionic lipids (e.g. cardiolipin, PIP_2_, etc.) with binding sites on integral membrane proteins.

To determine PMFs we first devised a scoring function that was based on the distribution densities of PIP_2_ around the binding site in order to find the most representative binding pose of the lipid. Steered MD simulations were used to generate a series of configurations along a direction defined by centre of masses of the receptor and the bound lipid, and umbrella sampling was used to estimate PMFs for the PIP_2_/receptor interactions. Comparing the PMFs revealed that PIP_2_ binding energetics at sites TM3/ICL2/TM4 and TM3/TM5 showed no significant difference between the inactive and active states of the receptor (Figure 5B and 5C). In contrast, for the TM1/TM2/TM4 and TM6/TM7 sites there was significantly stronger binding of PIP_2_ to the receptor in active state than to that in inactive state, especially for the TM6/TM7 site at which an increase of ~23 kJ/mol was observed (Figure 5D). This increase in PIP_2_ binding strength at TM6/TM7 is primarily due to that outward movement of TM6 which opens the intracellular side of the receptor and consequently allows PIP_2_ to bind more deeply and hence more tightly in this site (Figure 5E). Thus, the ingress of the anionic PIP_2_ molecules in the space between TM6 and TM7 may stabilize the outward movement of TM6 that is required for GPCR activation and G protein association, as has been suggested recently for other lipids (Dawaliby et al., 2015). Comparable phenomena have been reported for other lipids by MD simulations in simpler lipid bilayers (Caliman et al., 2017; Neale et al., 2015). However, in our simulations using an *in vivo*-mimetic membrane, the opening between TM6 and TM7 was almost exclusively occupied by PIP_2_, the multivalent anionic headgroup of which forms tighter interactions with the receptor than would be the case for other anionic phospholipids in the lower leaflet of the membrane, e.g. PS. To test this hypothesis, we carried out simulations on the receptor in active state and active + mini Gs state in a complex membrane devoid of PIP_2_ (SI Table S3), and calculated the PMFs for protein/PS interactions. The binding sites of PS on the receptor overlapped well with those of PIP_2_ (SI Figure S7). However, the interaction duration of PS was about one magnitude smaller than that of PIP_2_. Calculating PMFs, the binding energy of PS to the receptor in the active state and in the active + mini Gs state at site TM6/TM7 were −8.0 kJ/mol and −8.3 kJ/mol respectively, i.e. ~40 kJ/mol and ~50 kJ/mol weaker than that of PIP_2_ binding to the same site for the corresponding two conformational states respectively (SI Figure S8).

**Figure 5.**
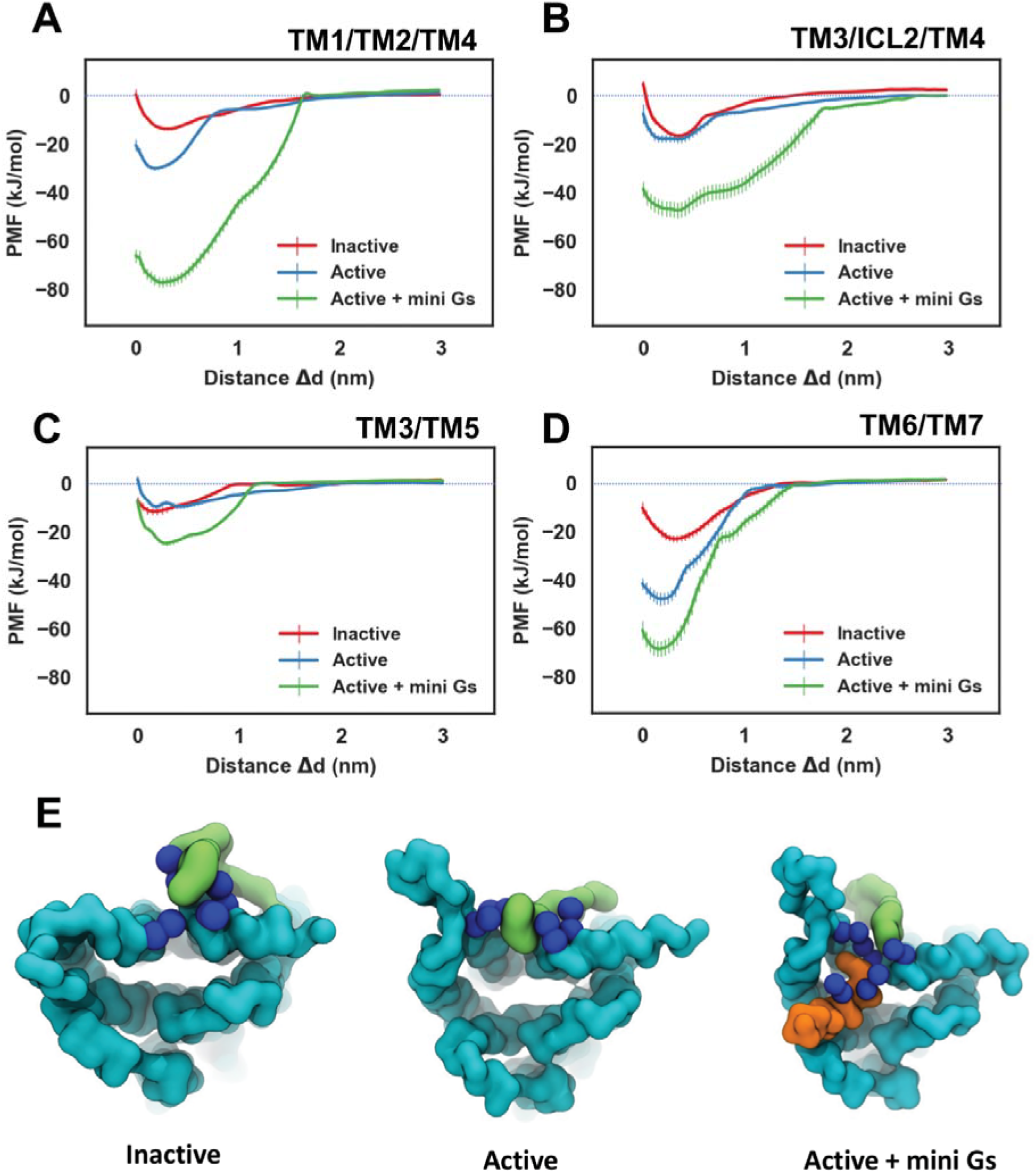
Potentials of mean force (PMFs) for PIP_2_ binding to the sites (see Figures 3 and 4) defined by TM1/TM2/TM4 (**A**), TM3/ICL2/TM4 (**B**), TM3/TM5 (**C**), and TM6/TM7 (**D**). The PMFs from the simulations of PIP_2_ bound to the inactive state, active state, and active + mini Gs state of the receptor are coloured in red, blue and green respectively. PIP_2_ bound to the TM6/TM7 site in the three conformational states is shown in (**E**) viewed from the intracellular side of the receptor. The receptor, the bound PIP_2_ molecule and the Gα α5 helix are coloured in cyan, green and orange respectively. The basic residues which form the binding site of TM6/TM7 (K233^6.35^, R29 1 ^7.56^, R293^8.48^, R296^8.51^) and from Gα α5 (R385, R389) are shown as blue spheres.

In the complex of A2aR and mini-Gs, the PIP_2_ binding sites are reinforced by adjacent basic residues of the mini-Gs protein, namely: R42 and R270 of mini-Gs near the TM1/TM2/TM4 site, K211 and K216 near the TM3/ICL2/TM4 site, R380 near the TM4/TM5 site, and R389 near the TM6/TM7 site (Figure 6). As the PIP_2_ molecule interacts with basic residues from both A2aR and the mini-Gs, it binds more strongly to all four sites, including the TM1/TM2/TM4 and TM6/TM7 sites which already showed state-dependency of the strength of interactions. Thus, PIP_2_ seems to both act a lipid bridge between the A2aR and the mini-G protein, and more importantly as a potential allosteric activator favouring the active + mini-Gs state of the receptor.

**Figure 6.**
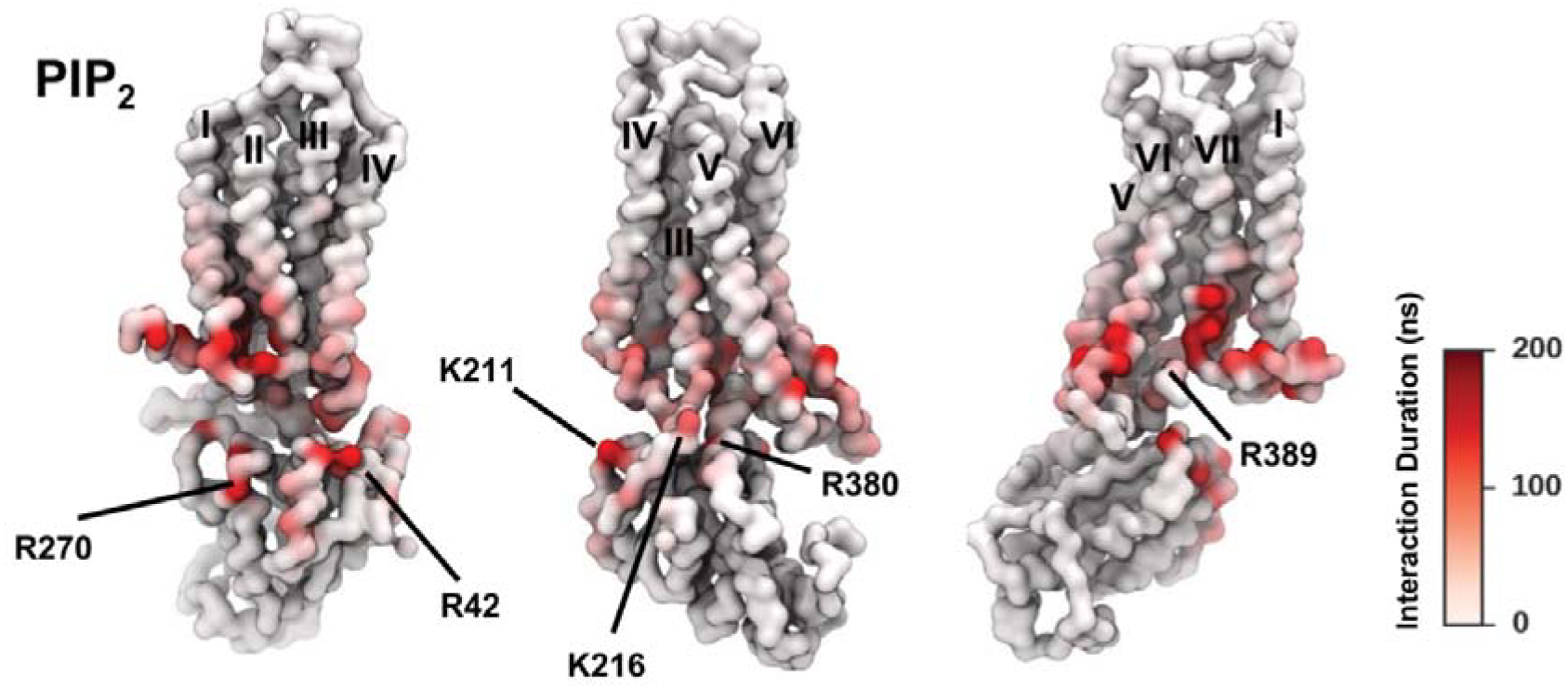
The duration of PIP_2_ interaction with A2aR in active + mini Gs state is mapped onto the receptor structure shown in three different orientations. Major interacting residues on mini Gs are labelled out.

### PIP_2_ enhances interactions of A2aR with mini-Gs protein

In the active + mini Gs state, we observed that the bound PIP_2_ molecules bridge the interaction between A2aR and mini Gs, which in turn suggests that PIP_2_ enhance the interaction between the receptor and the mini-G protein. To test this hypothesis, we calculated PMFs for the interaction energy between the A2aR and mini-Gs in the presence and in the absence of PIP_2_ (Figure 7). Simulation systems of the A2aR-mini-Gs-PIP_2_ complex in the complex membrane bilayer were generated via positioning mini Gs back to receptor structures in the active state simulations based on alignment with the crystal structure (PDB code 5G53). Independent PMF calculations were carried out on the three generated systems wherein the A2aR-mini-Gs complex showed lowest RMSD to the reference. Steered MD was used to generate a series of initial configurations along the membrane normal and umbrella sampling was used to calculate the PMF of A2aR-mini Gs interaction. The three independent repeats, albeit corresponding to slightly different initial A2aR-mini-Gs-PIP_2_ complex system, revealed similar sequences of events during the dissociation of A2aR and mini Gs. In this dissociation process, interactions between the mini-Gs protein and PIP_2_ molecules at sites at TM3/ICL2/TM4 and TM3/TM5 exhibited the most persistence to the pulling force. As illustrated by one of the repeats wherein R385, R380, and R373 on the Cα5 helix of mini Gs interacted with the PIP_2_ molecule bound to TM3/TM5 (PIP_2_ #2 in Figure 7B), and R42 and K216 on the β-strands of mini Gs interacted with the PIP_2_ bound to TM3/ICL2/TM4 (PIP_2_ #1 in Figure 7B), the interaction between K216 of mini Gs and the bound PIP_2_ held the mini Gs in contact until a break at ~42 ns when full dissociation occurred (Figure 7C). Regardless of the differences in initial configurations, the three independent PMF calculations yielded a consistent mini-Gs binding energy ~150 kJ/mol in the presence of bound PIP_2_.

**Figure 7.**
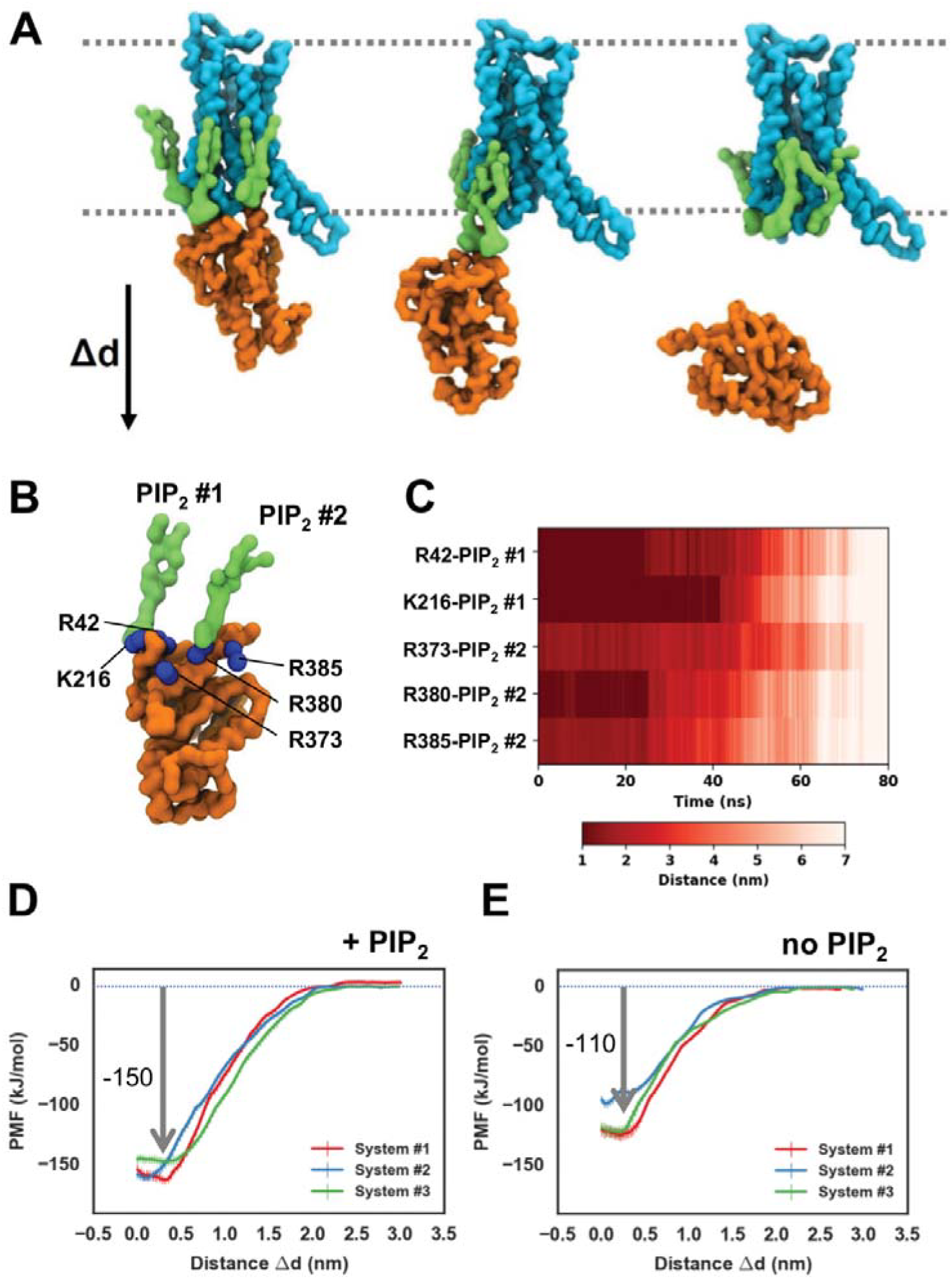
(**A**) An illustration of steered MD simulations pulling away the mini Gs from the A2aR along the z axis. The A2aR, the bound PIP_2_ molecules, and mini Gs are coloured cyan, green, and orange respectively. (**B**) The two bound PIP_2_s interact with basic residues on mini Gs, including R42, K216, R373, R380 and R385. (**C**) The distances between the two bound PIP_2_s and their corresponding contacting basic residues in the steered MD simulations. Potential of Mean Forces (PMFs) of A2aR-mini Gs association in the PIP_2_-containing membranes (**D**) and PIP_2_-deprived membranes (**E**). PMFs were calculated from three different systems and coloured differently for each membrane condition. Error bars represent the statistical error calculated by Bayesian bootstrap.

We then repeated the PMF calculation in the absence of PIP_2_. The initial configurations for Steered MD were generated from simulations of A2aR in the active state embedded in a complex membrane devoid of PIP_2_ using the same protocols. This resulted in a reduction of the free energy of interaction between the receptor and mini-Gs of ca. 40 kJ/mol compared to in the presence of PIP_2_ (Figure 7D and 7E). This reduction suggests a specific effect of PIP_2_ in stabilizing the receptor/G protein interaction, which could be explained from a structural perspective: PIP_2_ has a bulky headgroup of a phosphorylated inositol that is able to reach to the lower rim of the intracellular side of A2aR and to the mini Gs whereas PS has a small headgroup of serine that is limited in reaching out to mini Gs.

## DISCUSSION

We have performed a CG-MD simulation study of the interactions with different species of lipid molecule of a prototypical GPCR, the A2a receptor, in three different conformational states whilst embedded in a complex *in vivo*-mimetic lipid membrane. Ten different lipid species were included in our membranes, mimicking the lipid composition of a mammalian plasma membrane in a model where the concentration of each lipid species was sufficiently high to generate statistically valid data of protein/lipid interactions. The combinations of a multi-component lipid composition and different receptor conformational states allowed us to study changes in the spatial distribution and interactions of lipids in response to the receptor conformational changes. In our simulations, GM3, PIP_2_ and cholesterols predominated in the first layer of lipids around the receptor (Figure 2A), and their interactions with the receptor showed a degree of sensitivity to the receptor conformations (Figure 3). The differences in lipids interaction duration seen between the conformational states of the receptor suggest that the lipid binding affinity at the interaction sites changes during receptor activation. One functional outcome of such state-dependent interactions is that lipids may regulate the local conformational dynamics of the receptor that would be critical for ligand binding and downstream signalling. For example, the ECL2 loop has been shown to modulate ligand recognition, selectivity and binding (Klco et al., 2005; Ragnarsson et al., 2015). Our simulations suggest that the ECL2 loop is likely to be more flexible in the active state due to the decreased interaction duration of GM3 at the two sites (TM3/TM4/ECL2 and TM4/ECL2/TM5) to which this loop contributes, thus facilitating the entry and/or exit of ligand and modulating the kinetics of ligand binding. This influence of glycolipids on ECL2 may provide a structural explanation for the observation that GPCRs exhibit different ligand efficacies in different cell lines (Kenakin, 2002).

A key finding from our simulations is that the polyanionic lipid PIP_2_ enhances the interaction between the A2aR and a mini Gs protein. PIP_2_ molecules bound to cationic intracellular rim on the A2aR form an extended anionic surface at the cytoplasmic face of the receptor and thus facilitate the recruitment of G protein via formation of bridging interactions with basic residues on Gα. In the steered MD simulations of A2aR-mini Gs dissociation, we observed that the most resilient interactions were between PIP_2_ bound at the TM3/ICL2/TM4 site and basic residues of Gα S1-3, e.g. R42 and K216. Structural comparison between the Gα in the closed state (GTPγS-bound) and open state (receptor-associated) shows that K216, which is located on the short turn connecting β2 and β3 of Gα, remains solvent accessible in both states. Thus these interactions of PIP_2_ could be a major stabilizer during the initial stages of GPCR-Gα association. They also may provide a structural explanation for the observation that β2/β3 of the Gα subunit, whilst suggested by earlier biochemical studies to interact with GPCRs (Chung, 2013), did not form direct contacts with the GPCR in e.g. the crystal structure (PDB code 3SN6) of the β2-adrenergic receptor-Gs complex.

Crystal structures together with biochemical studies have revealed that the α5 helix of the Gα subunit undergoes rotational and translational movements during its activation by GPCR binding (Chung et al., 2011; Oldham et al., 2006; Rasmussen et al., 2011a). MD simulation studies suggest that the energy barrier between the inactive and active states of α5 is large, and the activated Gαsβγ in complex with GPCR is not stable when the receptor complex is embedded in a simple POPC membrane bilayer (Dror et al., 2015; Mnpotra et al., 2014). Based on our simulation data, we suggest that PIP_2_ bound to the TM3/TM5 site may facilitate the movements that α5 experiences during activation and help to stabilize the activated conformation. In our simulations, PIP_2_ showed similar affinities for the inactive state and active state when binding at the TM3/TM5 site, albeit with different interaction fingerprints. In the inactive state, the bound PIP_2_ had closer contacts with TM5, whereas in the active state the predominant contacts shifted towards TM3 (Figure 3C). In the active + mini Gs state, the bound PIP_2_ molecule moved further towards TM3 so that tight interactions were formed between PIP_2_ and basic residues from both the cytoplasmic end of TM3 and the α5 helix of Gα. Superimposing an inactive Gα protein (PDB code 1GOT) onto the model A2aR-PIP_2_-miniG complex based on the Ras-Homology Domain of Gα showed that the PIP_2_ molecule bound to the TM3/TM5 site from the active state simulations would interact with a basic residue (K341 in structure 1GOT) at the C-terminus of the α5 helix (Figure 8A). In contrast, superimposing an *active* Gα protein (PDB code 3SN6) onto the model A2aR-PIP_2_-miniG complex showed that the bound PIP_2_ from the active A2aR + mini Gs state simulations would interact with a basic residue in the middle of the α5 helix (R380 in structure 3SN6) (Figure 8B). Thus, by moving towards TM3 and sliding down the “basic ladder” on the α5 helix of the G protein, the bound PIP_2_ could help to draw the α5 helix into the binding pocket formed by the TM helix bundle of the GPCR and thus activate the G protein. Sequence alignment shows that the basic residues near the C terminus of α5 are conserve (Figure 8C), which suggests that this mechanism of PIP_2_-induced Gα activation may be a shared mechanism across different types of Gα. As to the selectivity toward different Gα the complementarity of the surface of Gα and that of the cytoplasmic side of the GPCR might play a major role (Baltoumas et al., 2013).

**Figure 8.**
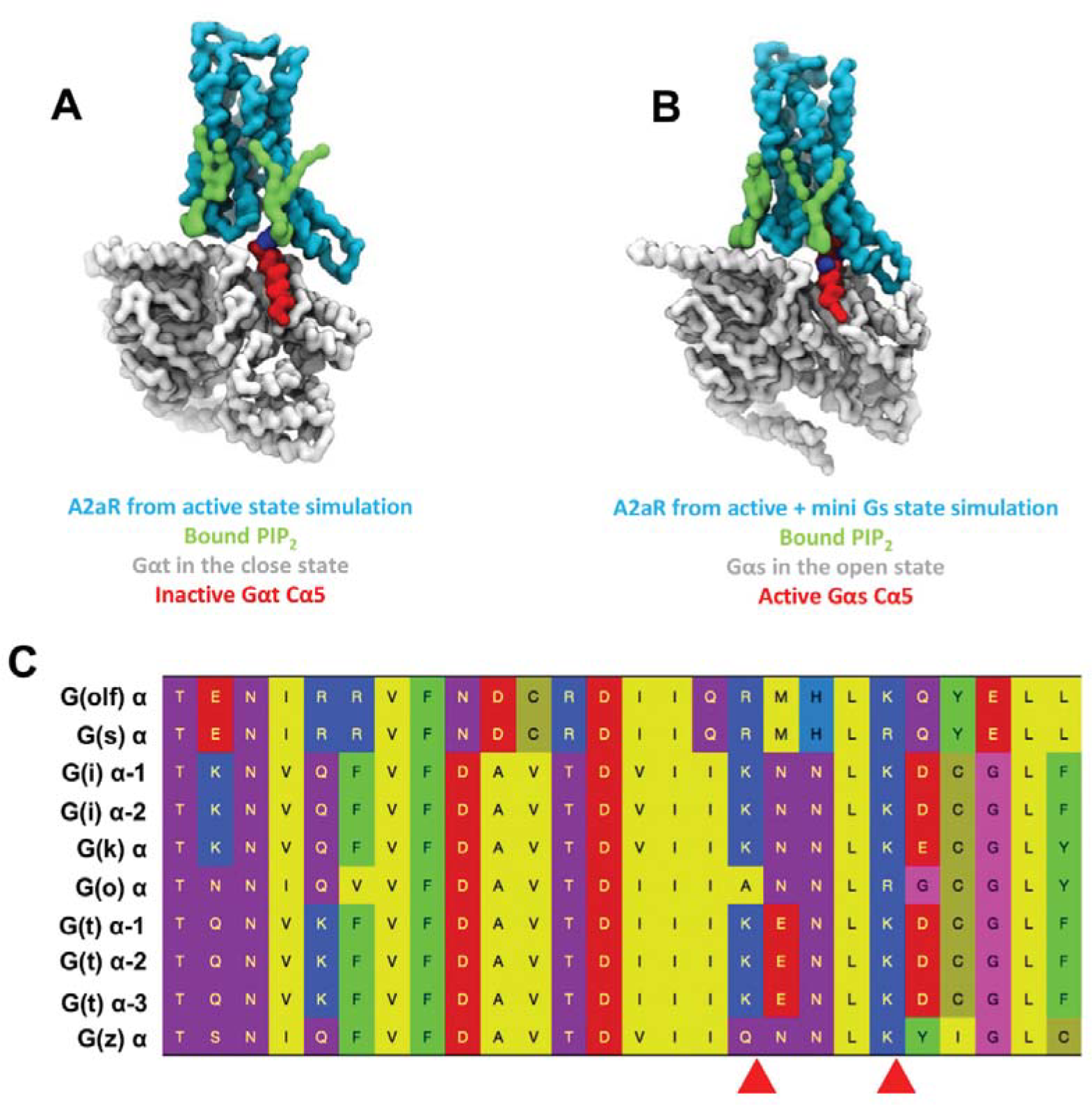
(**A**) Superimposition of the inactive Gαt (PDB code 1GOT) onto A2aR from the simulation of active state and the bound PIP_2_ molecules. The receptor and the bound PIP_2_ molecules are coloured cyan and green respectively. The α5 helix is coloured red while the rest of the Gαt protein is in grey. The basic residue in contact with the bound PIP_2_ molecule (K341) is shown as blue spheres. (**B**) Superimposition of the inactive Gαs (PDB code 3SN6) onto A2aR from the simulation of active state and the bound PIP_2_s. The α-helical domain of Gs (K88-V202) is omitted for clarity. The receptor and the bound PIP_2_ molecules are coloured cyan and green respectively. The α5 helix is coloured red while the rest of Gs in grey. The basic residue in contact with the bound PIP_2_ (R380) is shown as blue spheres. (**C**) Sequence alignment of α5 from different types of Gα. The conserved basic residues at the C terminal end are indicated by the red arrowheads.

In simulations of both the active state and the active + mini Gs state of the A2AR we observed stable binding of PIP_2_ at the site formed by TM6/TM7 (Figure 3C). We propose that such interactions may favour the outward movement of the cytoplasmic half of TM6 that is characteristic of GPCR activation. A similar stabilizing effect may be achieved by PS binding at the same site as revealed by our simulations in the absence of PIP_2_. Other phospholipids, including PG and PC, were reported to bind to this opening in atomistic simulations, resulting in different fingerprints of interhelical movements at the intracellular side of the receptor depending on the lipid headgroup properties (Neale et al., 2015). Such distinct conformational responses to different species of lipids could lead to differentiation of e.g. downstream signalling partners(Rose et al., 2014). This in turn suggests that the various micro-domains of plasma membranes may regulate GPCR functions via direct protein-lipid interactions (Dawaliby et al., 2015).

The possible interplay of different lipid species on protein-lipid interactions has been somewhat neglected in previous simulations of GPCRs. In our simulations, interplay was observed between different lipid species interacting with the same (or overlapping) sites. This is seen most clearly for PIP_2_ and cholesterol, which share a number of binding sites. The binding of cholesterol molecules at the TM1/TM2/TM4 and TM3/ICL2/TM4 sites was increased by synergistic interactions with PIP_2_, whereas cholesterol interactions at the TM6/TM7 site were decreased by competing interactions of PIP_2_ (Figure 3B and 3C). This phenomenon of lipid interplay will increase the complexity of the effects of bilayer environment on GPCR-lipid interactions.

Overall, our simulations support the likely modulatory role of the effects of membrane lipids (Dawaliby et al., 2015) on the conformational dynamics and hence functions of GPCRs. In particular, PIP_2_ is shown to have multifaceted effects on A2aR: it can stabilize the active conformation; enhance A2aR-mini Gs association; and may also aid the activation of the G protein. Lipid interplays were revealed by the usage of complex multi-component lipid bilayers, emphasizing the importance of membrane composition on modulatory effects on receptors. Our results highlight the integration of lipids with membrane receptors and suggest the existence of ‘mega-receptors’ the function and dynamics of which are governed by both the protein receptor and its bound lipids. This opens up new prospects for the pharmacology of GPCRs as their druggable space is expanded to include the bound lipids. The sensitivity of protein-lipid interactions toward the receptor conformational state and the lipid environment may thus provide a platform for designing subtype-selective and cell type-selective drugs.

## METHODS

### System setup

The inactive conformation of the A2a receptor was taken from the crystal structure 3EML (PDB code). The ligand and T4 lysozyme were removed and missing residues between P149 and H155, and between K209 and A221 were modelled using Modeller 9v9(Sali et al., 1995). The active state and the active + mini Gs state were both taken from the crystal structure 5G53 with the coordinates of mini Gs removed or retained respectively. Chain A of the receptor and chain C of the mini Gs were used. Missing residues of the receptor between G147 and G158, and E212 and S223 were modelled, while those in the mini Gs were discarded. Protein structure coordinates were converted to coarse-grained MARTINI representations using the martinize script(de Jong et al., 2013). Their secondary and tertiary structures were constrained using the ElNeDyn elastic network(Periole et al., 2009) with a force constant of 500 kJ/mol/nm^2^ and a cut off of 1.5 nm. The CG protein coordinates were then positioned in the centre of a simulation box of size 17×17×18 nm^3^ with its principal transmembrane axis aligned parallel to the z axis and embedded in a complex asymmetric membrane bilayer comprised of 10 lipid species using the insane script(Wassenaar et al., 2015). The membrane bilayer contained POPC (20%): DOPC (20%): POPE (5%): DOPE (5%): Sph (15%): GM3 (10%): Chol (25%) within the upper leaflet, and POPC (5%): DOPC (5%): POPE (20%): DOPE (20%): POPS (8%): DOPS (7%): PIP_2_ (10%): Chol (25%) within the lower leaflet (SI Table S1). To study the influence of PIP_2_ on A2aR-mini Gs association, we also ran simulations on the active state and active + mini Gs state in complex membranes deprived of PIP_2_. In these simulations, the lipid composition of the upper leaflet remained unchanged while the lipid concentrations of POPC, DOPC, POPE and DOPE in the lower leaflet were increased by 2.5% each (SI Table S2). 0.15 M NaCl was added to neutralize the system.

### Simulation parameters

The Martini coarse-grained force field version 2.2(de Jong et al., 2013) was used for protein and version 2.0 for lipids. All the simulations were performed using Gromacs 4.6(Abraham et al., 2015). The non-biased simulations were run in the isothermal-isobaric (NPT) ensemble equilibrium simulations. The temperature was controlled at 323 K using the V-rescale thermostat(Bussi et al., 2007) with a coupling constant of τ_t_ = 1.0 ps. The pressure was semi-isotropically controlled (i.e. independently in the *xy* plane and z axis direction) by a Parrinello-Rahman barostat(Martonak et al., 2003) at a reference ofp = 1 bar with a coupling constant of τ_t_ = 12.0 ps and compressibility of 3 x 10^−4^ Non-bonded interactions were used in their shifted form with electrostatic interactions shifted to zero in the range of 0–1.1 nm and Lennard-Jones interaction shifted to zero in the range of 0.9–1.1 nm. A time step of 20 fs was used with neighbour lists updated every 10 steps. Periodic boundary condition was used in x, y and z axis. For each conformational state, i.e. the inactive state, the active state and the active + mini Gs state, 10 simulation systems were independently constructed such that different random initial lipid configurations around the receptor were generated for every system. For the active state and the active + mini Gs state in PIP_2_-deprived systems, 2 independent simulation systems were generated for each state. 8 μs data were collected for all equilibrium simulation trajectories. An overview of the equilibrium simulations is listed in SI Table S3.

### Potential of Mean Forces calculations

We identified from the equilibrium simulations four PIP_2_ binding sites at the intracellular rim of A2aR. We then determined the potential of mean forces (PMFs) of PIP_2_ binding to these identified sites. To find the most stably bound PIP_2_ conformation, i.e. the conformation with the highest probability, we constructed for each binding site separately a scoring function based on the distribution density of each bead of the bound PIP_2_ the centre of mass of which were within 1.0 nm radius of all the basic residues in that binding site. All the PIP_2_ bound conformations were ranked according to the sum of beads’ scores, and the system snapshot that contained PIP_2_ bound conformation with the highest score was taken out. For generating the configurations of umbrella samplings, the Steered MD (SMD) simulations were carried out on the identified bound conformations *in situ*, i.e. in the lipid environment from the non-biased equilibrium simulations. The bound PIP_2_ molecules were pulled away from the receptor in the membrane plane in a direction defined by the vector between the centres of mass (COMs) of the receptor and of the bound lipid. A rate of 0.05 nm/ns and a force constant of 1000 kJ/mol/nm^2^ was used. The starting configurations of the umbrella sampling were extracted from the SMD trajectories spacing 0.05 nm apart along the reaction coordinate. 50 umbrella sampling windows were generated, and each was subjected to 1.5 μs MD simulation, in which a harmonic restrain of 1000 kJ/mol/nm^2^ was imposed on the distance between the COMs of the receptor and the bound lipid to maintain the separation of the two. The PMF was extracted from the umbrella sampling using the Weighted Histogram Analysis Method (WHAM) provided by the GROMACS *g_wham* tool(Hub et al., 2010). A Bayesian bootstrap was used to estimate the statistical error of the energy profile, which is shown as error bars in Figures 5A-5D.

To study the impact of PIP_2_ on A2aR-mini Gs association, we calculated the PMFs of this association in two membrane bilayers, i.e. one with 10 % of PIP_2_ in the lower leaflet, and the other without PIP_2_. To mimic the association process in physiological condition, we generated the A2aR-mini Gs complex structures via putting the mini Gs back to the A2aR structure from the non-biased equilibrium simulations of the active state that showed lowest RMSD based on the complex crystal structure 5G53 (PDB code). Again, this process was carried out *in situ*, i.e. with the membrane bilayer from the non-biased equilibrium simulations. Three systems were generated independently for the PIP_2_-containing and PIP_2_-deprived simulations respectively. In the steered MD simulation, the mini-Gs was pulled away from the receptor along z axis (normal to the membrane plane) at a rate of 0.05 nm/ns using a force constant of 1000 kJ/mol/nm^2^. The distance between the COMs of the receptor and the mini-Gs was defined as the 1D reaction coordinate and the pulling processed covered a distance of 3 nm. Similar protocols as used in the PMF calculation of PIP_2_ binding were followed. 50 windows were generated by extracting configurations spacing 0.05 nm apart along the reaction coordinate. Each window was subjected to 1 μs of simulations with a harmonic restrain of 1000 kJ/mol/nm^2^ imposed on the reaction coordinate. WHAM was used to calculate the PMF from umbrella sampling. Statistic errors were calculated by the Bayesian bootstrap which are shown as error bars in Figures 6D and 6E. An overview of the PMF calculation simulations is listed in SI Table S4.

### Analysis

The radial distribution functions (RDFs) were calculated as the distribution of the centre of mass of lipid molecules to the surface of the receptor via Gromacs tool *g_rdf*. The area per lipid (APL) was calculated using Voronoi tessellation provided by the python Scipy package (version 0.19.1). Phospholipids were represented by the midpoint of GL1 and GL2 beads, i.e. the two beads representing the glycerol group; Sphingolipids were represented by the midpoint of AM1 and AM2 beads, i.e. the two beads representing the sphingosine head group; Cholesterols were represented by the ROH bead, i.e. the hydroxyl group. The tessellations at simulation box boundaries and adjoining the receptor were calculated taking into account the periodic boundary conditions and the position of beads from the receptor, respectively.

The *k*_*off*_ values for bound lipids were estimated by curve-fitting to the decay of interaction durations as a function of time. The interaction durations of the lipid species of study to a given residue were collected from the 10 equilibrium simulations of each receptor conformational state. A distribution density function was calculated from these interaction durations and was then fitted to a mono-exponential curve of *N* = *Ae*^−*k_off_t*^.

In estimating lipid interaction durations, a dual-cut-off strategy was adopted. Continuous lipid binding to a given residue was defined as starting when the centre of masse (COM) of the lipid was closer than 0.55 nm to that of the amino acid residue, and as ending when the COM of the lipid moved more than 1.4 nm away from that of the residue. The duration between these two events was taken as the lipid interaction duration with a given residue.

Protein-lipid interaction of concurrently bound lipids can operate in either a synergistic or a competing fashion (see Results). To quantify this effect, we calculated the Pearson’s correlation coefficient (P.C.C) of interaction duration of two cohabiting lipid species. By definition, 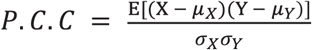 where X is the sample values of the interaction duration of lipid Y, *μ*_*X*_ and *μ*_*Y*_ is the sample values of the interaction duration of lipid Y, *μ*_*X*_ and *μ*_*Y*_ are the averages of sample X and sample Y respectively, and *σ*_*X*_ and *σ*_*Y*_ are the standard deviations of sample X and sample Y. E is the expectation. Thus, the P.C.C of interaction duration was calculated as 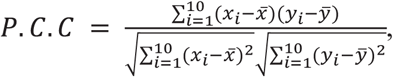, where *x*_*i*_ and *y̅* were the average interaction durations of the two lipid species of study in simulation repeat *i*, while *x̅* and *y̅* were the average interaction durations of all the simulation repeats.

## ACKNOWLEGEMENTS

Research in M.S.P.S. group is supported by: Wellcome (208361/Z/17/Z), BBSRC (BB/R00126X/1), and PRACE (Partnership for Advanced Computing in Europe; 2016163984). CVR acknowledges an ERC Advanced Grant ENABLE (641317), an MRC Programme Grant (G1000819) and a Wellcome Trust Investigator Award (104633/Z/14/Z). W.S. acknowledges support from the Newton International Fellowship. This project made use of time on ARCHER via the HECBioSim, supported by EPSRC (EP/L000253/1).

